# The Universal Endurance Microbiome?

**DOI:** 10.1101/2022.07.20.500882

**Authors:** Hope Olbricht, Kaitlyn Twadell, Brody Sandel, Craig Stephens, Justen Whittall

## Abstract

Billions of microbial cells sculpt the gut ecosystem, playing essential roles in human physiology. Since endurance athletes’ performance is often physiology-limited, understanding the composition and interactions within these athletes’ gut microbiomes could lead to improved performance. Previous studies describe differences in the relative abundance of bacterial taxa when comparing athletes versus controls or athletes before and after an endurance event, suggesting the existence of an “endurance microbiome”. However, there are inconsistencies among studies in which taxa correlate with extended physical exertion. Although these studies employed similar barcoding methods, variation in downstream bioinformatic analyses makes it difficult to determine whether inconsistencies are due to methodological differences or biological factors. Herein, we created a metagenomic bioinformatics workflow reanalyzing four 16S rDNA sequence datasets reflecting endurance athletes’ gut microbiomes, looking at alpha diversity, changes in relative abundance of gut microbiome genera, changes in pairwise correlations between bacterial genera and compared bacterial association networks. There were no significant differences in alpha diversity between any of the four treatment group comparisons. For relative abundance, there were no consistent differences in all four datasets, and only two genera were significantly different in 50% of the datasets. Although many genera showed changes in pairwise correlations in endurance microbiome samples from individual datasets, none were consistent across datasets. Collectively, these results suggest that either there is no universal endurance microbiome, or that it remains elusive even after controlling for the bioinformatic workflow and statistical analyses. Using this data, a power analysis indicates that sample sizes 150- to 800-fold larger than these published studies would be necessary to detect a 10% difference in relative abundance. Furthermore, 10- to 20-fold more samples will be needed to control for the multitude of covariates (genetic, metabolic, dietary, environmental, and pharmacological factors) that mold the gut microbiome of athletes and non-athletes alike.

*I’m going to work so that it’s a pure guts race at the end, and if it is, I am the only one who can win it. -* Steve Prefontaine

## Introduction

Endurance has played an important role in human history from our origins in the African savannah, through historical times (i.e. the origin of the marathon) to the more recent fascination with ultra- endurance events. Success in endurance events is assumed to be a product of training, genetics, and psychological preparation in order to withstand extreme mental and physical challenges [1]. Physiologically, the factors limiting performance in endurance events have traditionally been divided into two broad categories - aerobic (i.e. VO_2_ max) and anaerobic (i.e. lactic acid) [2]. However, what if one of the limiting factors to success in endurance events wasn’t human at all?

We are as microbial as we are human. Bacterial cells associated with the human body are more common than human cells (∼100 trillion) [3]. More than 1000 bacterial species may be found in the intestines of each person, and over 70% of the total human microbiome is contained in the gut [4]. Whereas human cells are slowly replaced with identical (or nearly identical) copies, the cells of the gut microbiome are constantly changing due to immigration, emigration and differential rates of division based on the dynamic gut environment, to which they are much more sensitive and responsive than human epithelial cells. If our fuel tanks are lined with bacteria, it is not surprising that the gut microbiome may affect performance in fuel-limited sports such as endurance events. Furthermore, the dichotomy between human physiology and microbial metabolism is increasingly difficult to disentangle, as more human genotype/microbiome interactions continue to be discovered [5].

In some studies, exercise has been associated with increased gut microbial diversity, increased *Bacteroidetes*-*Firmicutes* ratio, and proliferation of bacteria which can modulate mucosal immunity and improve barrier functions. All of these changes could contribute to increased performance, decreased inflammation, decreased gastrointestinal (GI) distress, and faster recovery times [6], as well as possible protection against GI infections [7–11]. However, factors other than intensive exercise could also contribute to these reported changes in the microbiomes of endurance athletes. In fact, more than 100 factors have been suggested to affect the composition of the gut microbiome, including (but not limited to) method of birth, diet, sex, age, antibiotic use, geographic region, stress level, and disease history [12]. For example, dietary behaviors like carbo-loading, a traditional prelude to an endurance event, could increase *Prevotella* independent of physical exertion [13]. Consumption of probiotics could also affect the microbiome (for example, see [14]). A recent review found mixed effects of probiotics on performance, as 17 studies showed no significant results of probiotic consumption and seven showed an improvement in performance (summarized in [13]). Probiotics may also protect against some upper respiratory tract infections which in athletes could result in longer stretches of continuous, intensive training [15]. Interactions among bacteria may be important, as multi-strain probiotics are more likely than single strains to lead to improved endurance performance [13].

The concept of an “endurance microbiome” suggests there are assemblages of gut bacteria that are more common in athletes compared to sedentary controls, or gut bacteria that are enriched after an endurance event compared to before the event. This hypothetical endurance microbiome could be characterized by changes at multiple levels such as (1) increased overall diversity, (2) different levels of abundance, and (3) the emergence of bacterial associations. Below, we briefly summarize these three metrics in relation to the endurance microbiome.

To date, studies on the effects of exercise on gut microbiome diversity have exhibited mixed results. Some investigations describe increased microbiome diversity in athletes [6, 16–20], while others report no significant change in diversity [21–23]. In regards to relative abundance, three studies have come to largely different conclusions. First, In a comparison of runners before and after the Boston Marathon, Scheiman et al. [24] found only *Veillonella* exhibited a significant increase in abundance after the marathon. The authors proposed that *Veillonella* could in fact improve endurance performance by limiting accumulation of metabolites (lactate/lactic acid) that limit success in an endurance event [24]. They further showed that lactic acid could cross the blood-gut barrier, and therefore could plausibly be a carbon source for the gut microbiome. Second, when Zhao et al. [23] examined Chinese half marathon runners, they found 12 genera whose abundance increased after the event, not including *Veillonella*. Third, Peterson et al. [18] compared cyclists with variable training intensities and found some of the same genera as Zhao et al. [23], yet identified several new genera with differential abundance based on metagenomic and transcriptome investigations. These studies identify many candidate groups for the endurance microbiome, but due to methodological differences in bioinformatics workflows, it is difficult to make direct comparisons among the studies.

The endurance microbiome could also manifest itself in associations among bacteria, or groups of bacteria. For example, several studies have highlighted a putative trade-off between two common groups of gut bacteria implicated in endurance athletes - *Prevotella* and *Bacteroides* [25, 26]. Other investigations have found strong correlations among bacterial lineages and in response to environmental stresses [27]. Many exercise-related studies have looked for inter-bacterial associations. *Prevotella* has been implicated in athletic performance and associated with *Streptococcus, Enterococcus, Desulfovibrio, Lachnospiraceae*, *Succinivibrio*, *Oscillospira*, *Xylanibacter,* and *Butyrivibrio* [20]. Nevertheless, Gorvitovskaia et al. (<u>26</u>) report no consistent bacterial correlations among the four studies they reviewed. Potentially, bacterial interactions in the endurance microbiome are broader than pairwise correlations - more representative as connectivity networks of many bacteria. The broader network of connectivity underlying the endurance microbiome is largely unexplored [28], and no meta-analysis of multiple endurance microbiomes have compared networks using the same methodology across datasets.

We hypothesize that if an endurance microbiome exists, it should manifest regardless of geographic location, type of sport, or specific endurance event. Our goal is to reanalyze several endurance microbiome studies using a single bioinformatics pipeline and consistent downstream statistical analyses in order to determine whether there are repeated changes in diversity, relative abundance, and/or associations between genera in response to intensive endurance exercise.

## Methods

### Datasets

After searching the literature for relevant studies (Supplemental Table 1), we selected and reanalyzed three gut microbiome datasets involving endurance athletes (Table 1). All were composed of 16S rDNA amplicon sequence data collected on the Illumina platform [18, 23, 24]. Below is a brief description of each.

**Table 1.**
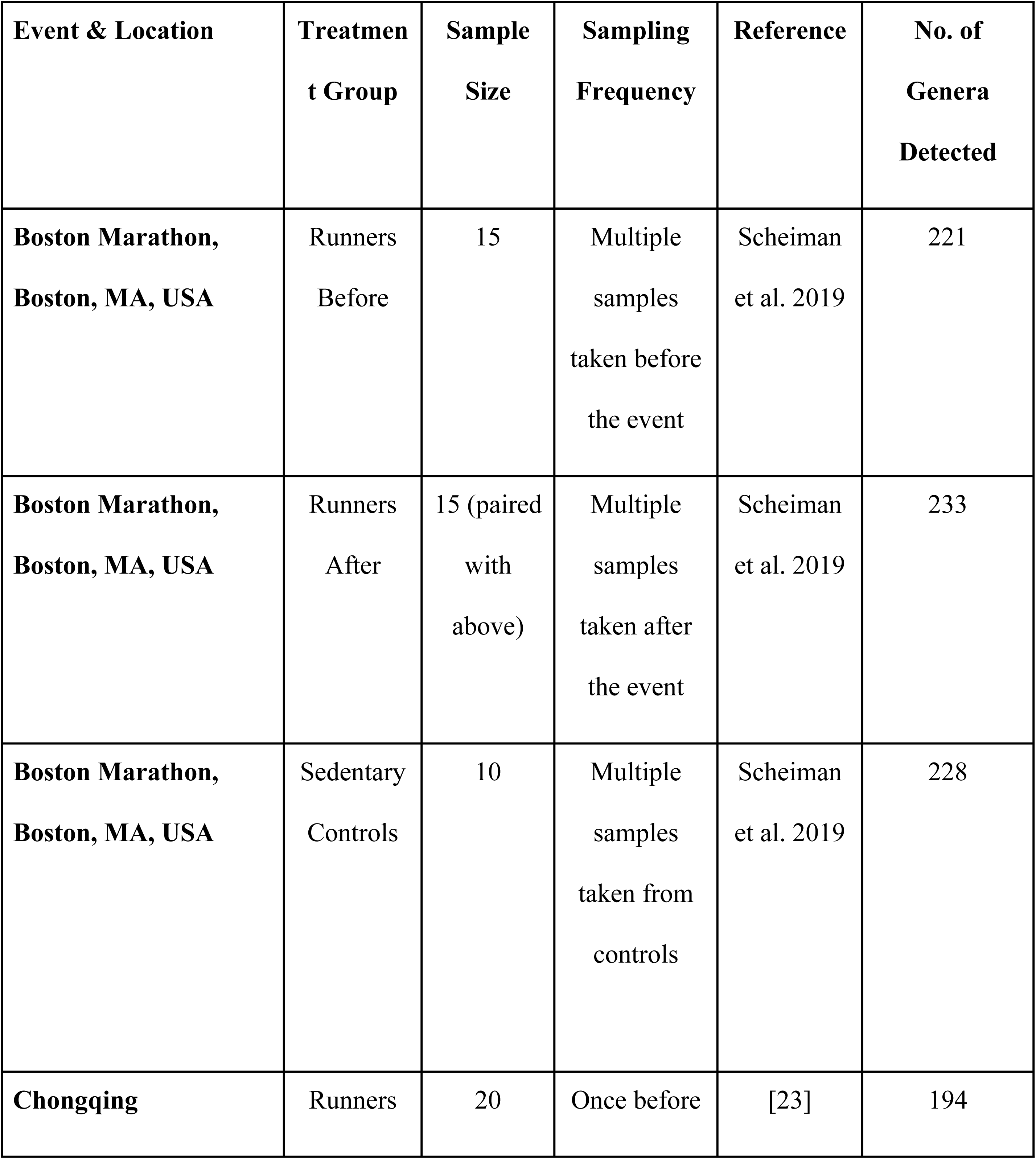
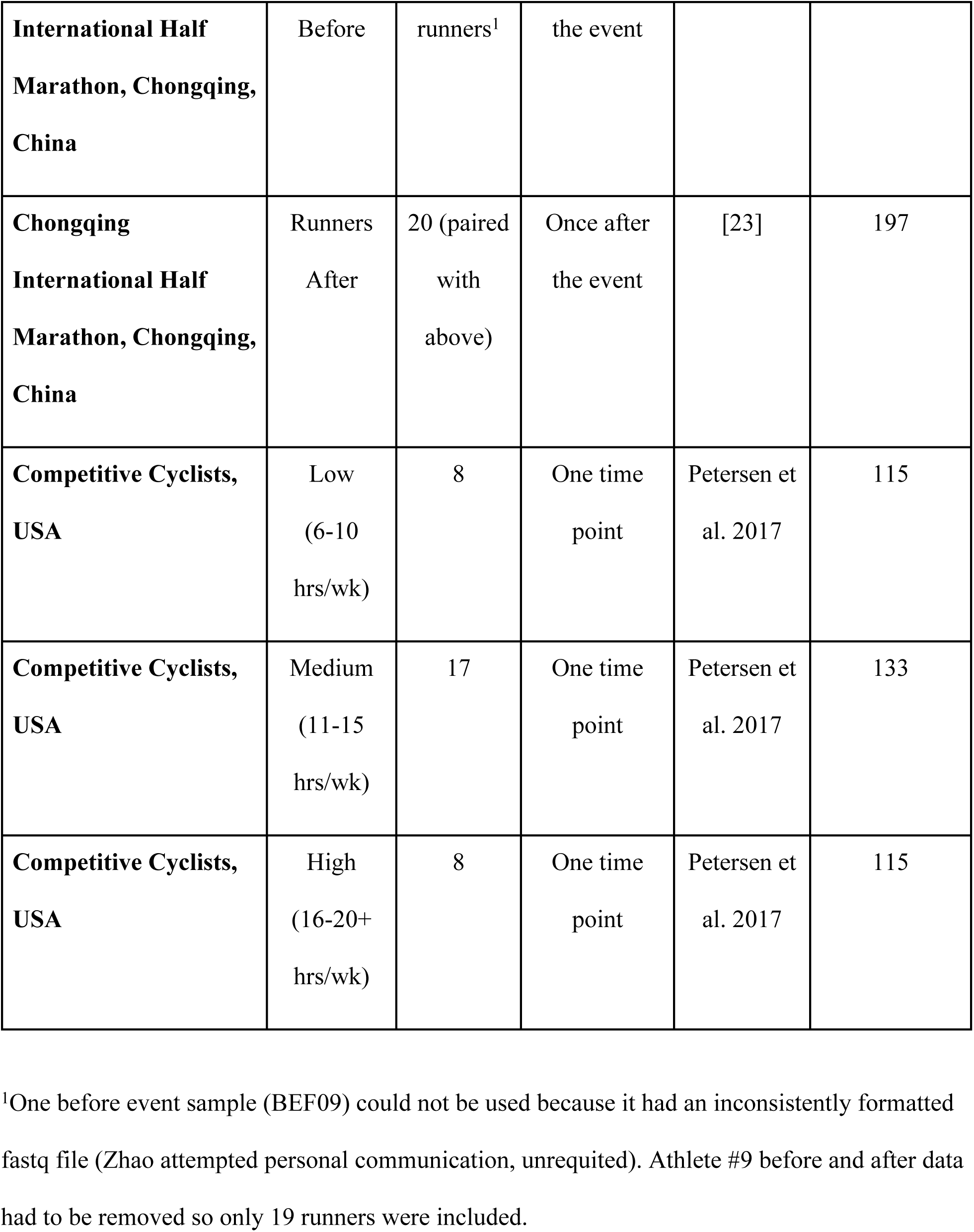
Treatment groups. Data reanalyzed in this work and number of microbial genera detected in our bioinformatics pipeline.

*Boston marathon study* - Scheiman et al. [24] recruited 15 elite athletes running in the 2015 Boston Marathon, along with 10 sedentary controls. They conducted amplicon sequencing on 209 fecal samples taken daily from participants up to one week before to one week after the marathon. 150 bp Illumina paired-end reads were processed with the DADA2 pipeline and phyloseq. Generalized linear mixed-effect models and leave-one-out cross validation were used to confirm significant associations. According to their Supplemental Data, some samples were “rerun” on the Illumina sequencer. In these cases, we only used the rerun data (i.e. SG10 and SG27).

*Chongqing half-marathon study* - Zhao and colleagues [23] recruited 20 amateur athletes who were running in the 2016 Chongqing International Half Marathon. A total of 40 fecal samples were collected - each runner was sampled the morning before the race, and again the after the race. Raw data consisting of 250 base pair paired-end reads were generated using an Illumina HiSeq of the fecal samples. Two samples from runner nine were eliminated from our reanalysis because of ambiguous labeling of the data. Each participant was given the same kind of food during the period between the first and second sample collection.

*Cyclist study* - Petersen et al. (<u>18</u>) studied 33 cyclists categorized into four non-overlapping training groups based on their average training time per week: 6-10 h, 11-15 h, 16-20 h and 20+ h per week. A total of 33 samples were collected, one from each cyclist. Either 125 or 150 bp paired end reads were generated using Illumina NextSeq and HiSeq. We grouped these into low (6-10; n = 8), medium (11-15; n = 17), and high (16-20+; n = 8) categories for diversity analyses, then focused on the two most extreme training groups (with balanced sample sizes) when searching for a universal endurance microbiome (low vs. high).

### Target genera

Among the Boston Marathon, Chongqing half-marathon and cyclist studies, there were 1, 12, and 6 bacterial genera identified as having significantly different abundances between treatment groups, respectively (Supplemental Table 2). Of these, no genus was identified in all three studies and only three genera were found in two studies. We used the 16 unique genera from all three studies as our “target genera” in a hypothesis testing framework (using alpha < 0.05 as a cutoff). Subsequently we expanded our analyses to all remaining genera, and corrected for multiple tests, since the comparisons were not based on *a priori* hypotheses.

The target “genera” from Zhao et al [23] included both individual species (e.g. *Prevotella corporis*) and genus-level groups. The authors emphasized the significantly differential abundance in the family Coriobacteriaceae before and after the half marathon, but we only included *Collinsella* since we assume this genus was driving the significant result (see their Fig. 2B). We did not include “unclassified Porphyromonadaceae’’. Finally, they report “*Phaseolus vulgaris*” in their Fig. 2 results [23], but they discuss *Romboutsia* later in their findings. We assume this is a technical mistake (*Phasaeolus vulgaris* being a flowering plant genus and not a bacterial genus). *Phaseolus* sensu Zhao et al. [23] is hereafter treated as *Romboutsia*.

### Microbiome assembly (bioinformatics pipeline)

Although there are many published tools for measuring bacterial abundance from 16S amplicon sequencing using the Illumina platform (QIIME [29]; MOTHUR [30]; DADA2 [31]), we developed a simple workflow in Geneious Prime 2021.2.2 by adapting the Amplicon Metagenomics tutorial (https://www.geneious.com/tutorials/metagenomic-analysis/). Raw data from published studies was downloaded from usegalaxy.org in the form of fastq files, and imported into Geneious Prime. We used Illumina paired-end inward pointing with sequences interlaced within each fastq file with an expected amplicon size of 292 base pairs. The minimum quality (q) cutoff was empirically determined to be 13 in a pilot study, and the merge rate was set at “very high” in order to maximize reads mapped, while reducing incorrectly mapped reads. After using BLAST to compare against the bacterial 16S database, sequences were classified to genus level based on 90% minimum overlap identity, per Geneious’ Amplicon Metagenomics tutorial & Sequence Classifier tutorials. (We recognize that 95% 16S identity has often been used as a cut- off for bacterial genus-level operational taxonomic units [32]; our more generous cut-off was intended to maximize the number of classified reads.) All subsequent analyses were based on the proportion of reads hitting a genus compared to total reads mapped per sample (per Gloor et al. [33]). Classifications were pruned to genus, with all higher taxonomic level BLAST results removed. Our Geneious workflow is available from FigShare (DOI: 10.6084/m9.figshare.19799203).

### Diversity

We measured alpha diversity for each dataset and each treatment group as the number of unique genera identified. We then used Simpson and Shannon indices to compare diversity in light of each genus’ relative abundance (vegan packages used). For the cyclist study [18], diversity measures among the three independent treatment groups were compared using an ANOVA in R. All other diversity comparisons for the Boston marathon and Chongqing half-marathoners were done using independent samples (athletes vs. controls) or paired (before vs. after) t-tests in R.

### Relative abundance comparisons

*Hypothesis-Driven Approach -* Datasets were examined for specific genera exhibiting significantly different abundances in athletes vs. controls or athletes before vs. after an event. These genera were then tested for significant differences in relative abundance for all three datasets for all four comparisons of treatment groups using the Wilcoxon test with continuity correction in R (alpha = 0.05). Since the Boston marathon study [24] had multiple samples collected before and after the event, the average relative abundance before and after was used, as well as averages for the multiple samples taken from the sedentary controls.

*Data Exploration -* To explore previously unreported differences in bacterial abundance among these datasets, relative abundances between treatment groups were examined for all remaining genera detected in the gut microbiomes (implemented in R library/package). We applied a Bonferroni correction for multiple testing (Boston marathon control vs. athletes = 282 tests; Boston marathon “athletes before” vs. “athletes after” = 282 tests; Chongqing half-marathon = 198 tests; Competitive cyclists = 148 tests).

### Changes in bacterial associations

*Correlated Abundances and Changes Therein -* Correlations in relative abundance between bacterial genera were tested for each treatment group individually. The analyses were restricted to the genera detected in at least 75% of all samples to avoid spurious correlations caused by large amounts of missing data (i.e. genera with no reads mapped). The Pearson’s correlation coefficient (r) and associated p-value were determined in R, then Bonferroni correction for multiple testing was applied (Boston marathon “athletes before”, “athletes after” & controls = 1596 tests; Chongqing Half-Marathon “athletes before” & “athletes after” = 3081 tests; Cycling “low”, “medium” & “high” = 861 tests). Significant correlations within treatment groups were then examined to see if they changed between treatment groups. Specifically, we examined changes in significant correlations between treatment groups for the three datasets to determine which associations between genera change in athletes vs. controls [24], before versus after [23, 24], or among training volume levels [18].

*Networks of Bacterial Associations (NetCoMi) -* Potential microbe–microbe association networks underlying the gut microbiomes of endurance athletes were explored using NetCoMi [28]. We looked for predictable changes in the connectivity of microbial association networks among treatment groups using quantitative comparisons across networks. Possible differences in the hierarchical clustering of bacterial interactions were examined between controls vs. athletes, athletes before vs. after and athletes involved in differing levels of training in the four datasets. Raw read counts from the Geneious bioinformatics pipeline were pruned to genera, then filtered for samples with 1000 reads or more and limited to the 100 genera with the highest number of reads [28]. We constructed the network with SparCC [34] which accounts for the compositional nature of the data by applying a centered log-ratio transformation (clr) [35, 36]. Furthermore, SparCC is less likely to identify spurious correlations compared to Pearson correlations [37]. Singular nodes were removed when present in one network and when comparing multiple networks.

A t-test was applied (alpha = 0.001 for Boston marathon samples [24]; alpha = 0.1 for Chongqing half-marathon samples [23]; alpha = 0.1 for the cycling study [18]) to reduce the network to a tractable size with a local false discovery adjustment [38] to select which edges to include in the network (“sparsification” [36]). To analyze the network, we applied the fast greedy modularity optimization algorithm for finding community structure (“cluster_fast_greedy” [39]) with hubs defined based on their eigenvalue (degree of connectedness to other nodes with high connectedness). A high eigenvector of centrality score means that a node is connected to many nodes who themselves have high eigenvector scores [40].

For each treatment group individually, we compared the relative size of the largest connected component of the resulting network (LCC = the connected component with highest number of nodes), betweenness centrality (the degree to which a node lies on paths between other nodes), closeness centrality (distance between a node and all other nodes), and degree centrality (# of edges = measure of co-occurrence) [41]. We also assessed dissimilarity (1 - edge weight) and average path length for each of the three treatment groups.

When comparing networks, the layout represents the optimal “union” of each pair of networks (athletes before vs. after; controls vs. athletes after; low vs. high training groups). Node size represents the centered log-ratio transformation of the number of reads per genus and node color distinguishes different clusters.

To compare two networks at a time (e.g. “athletes before vs. after” and “controls vs. athletes”), we examined network size (LCC), positive edge percentage (positively biased), hub taxa, adjusted Rand index [35], and the top 10 taxa showing the largest differences in three centrality measures (degree, betweenness and closeness).

We conducted several statistical comparisons for these pairs of networks (“athletes before” vs. “athletes after” and controls vs. “athletes after”) using CompareNet in NetCoMi [28]. To test whether the two similarity matrices are significantly different from one another, we examined Jaccard indices for degree, betweenness, closeness, eigenvector and hub taxa. A Jaccard index equal to zero indicates completely different matrices. In addition, we used the adjusted Rand index (ARI) which measures the similarity between clusterings (ARI = 0 means random clusterings in the two networks being compared; ARI = 1 means perfect agreement between clusterings in the two networks being compared) [42]. For Rand index, a p-value below 0.05 means that ARI is significantly higher than expected for two random clusterings. Significance for the Rand index was determined from 1000 permutations. We also used permutation tests (n = 1000) to determine if any genera showed significant differences in their degree, betweenness and closeness when comparing networks. Briefly, to generate a null distribution, the treatment group labels were randomly reassigned to the samples keeping the group sizes constant. The network metrics are then re-estimated for each permutation. We applied the local false discovery rate correction (lfdr) to correct for multiple testing.

We identified differences between the networks using Diffnet in NetCoMi [28]. To assess significantly different associations in the network, we applied Fisher’s Z-test after adjusting the p-value using the local false discovery rate. Occasionally, this produced an empty network (e.g. Scheiman “athletes before” vs. “athletes after”) in which case we loosened the filter in NetConstruct to ensure sufficient numbers of overlapping bacterial genera were able to be compared during the Diffnet analysis.

All R scripts for all post-Geneious data analyses are available from FigShare (DOI: 10.6084/m9.figshare.c.6036347).

## Results

The bioinformatics pipeline assembled and mapped the majority of the raw reads. For the Boston marathon and cyclist samples, ∼60% of reads mapped to a bacterial genus. (The remaining 40% did not meet the 90% identity threshold for known genera, and were removed). For the half- marathoners, ∼80% of reads mapped, reflecting the longer read lengths and thus increased likelihood of accurately mapping to a single genus.

### Diversity

Overall diversity was measured as the total number of genera detected (richness) and using Simpson and Shannon’s diversity indices. Variation in the number of genera detected was much higher among datasets than between treatment groups within each dataset (Table 1). We found 280 genera among the three Boston marathon treatment groups, 197 genera between the two Chongqing half-marathon treatment groups, and 148 genera among the three cyclist training groups.

In all datasets, a small number of genera make up the bulk of the microbiome (Fig 1). There was some overlap among studies. For example, *Faecalibacterium* was consistently one of three to five genera that comprise greater than 50% of the microbiome in treatment groups across datasets. In addition, *Bacteroides* and *Blautia* are very abundant in most treatment groups. However, there were also numerous differences. The most easily observable richness difference concerned the genus *Prevotella*, which was much more common in the medium and high cyclist training groups versus the low training group (Fig 1C), and was also rare in runners from the other datasets.

**Figure 1.**
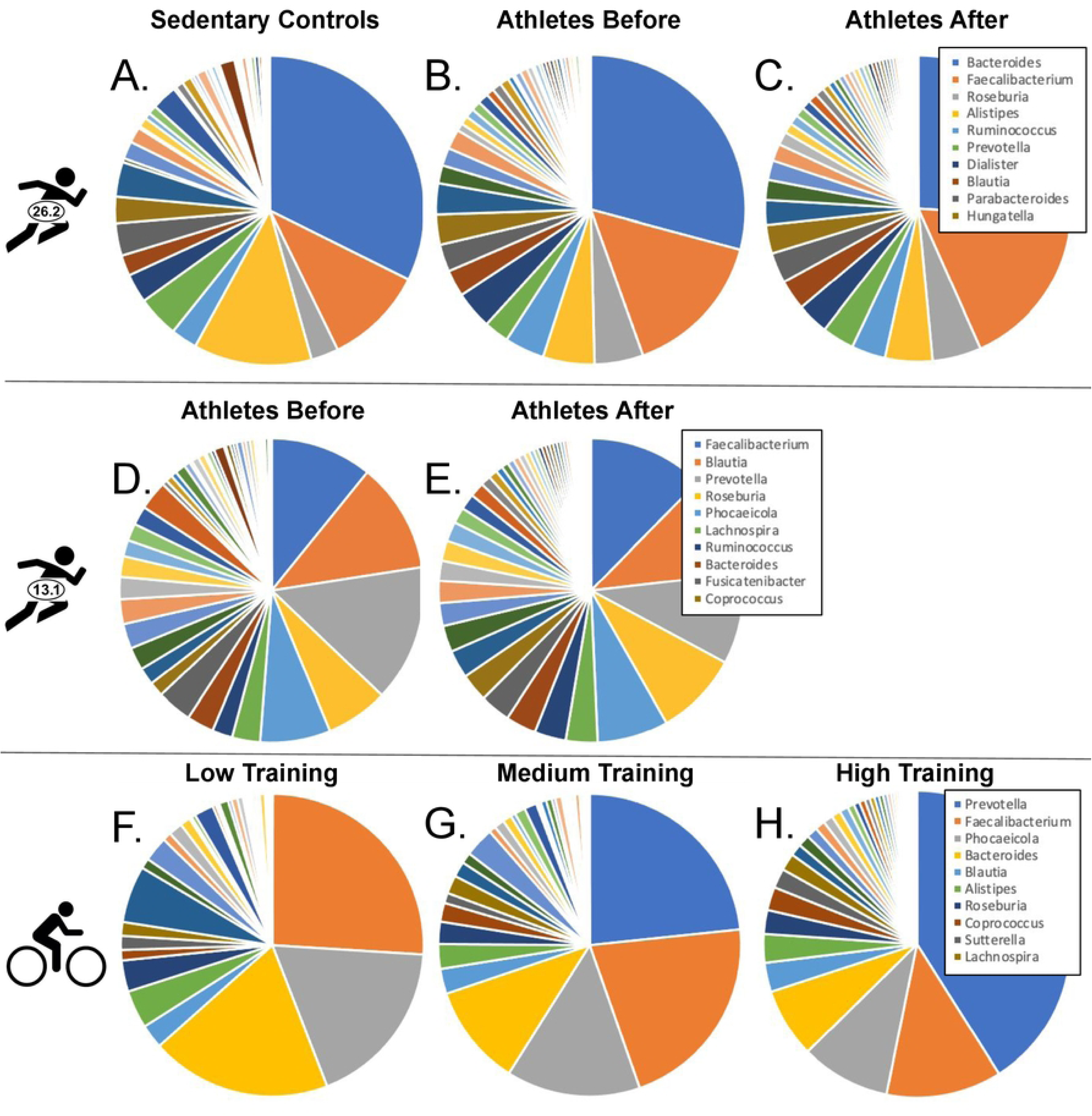
Average proportions of bacterial genera in the eight comparison groups. Data are from Boston Marathoners (A); Chongqing Half-Marathoners (B); professional cyclists (C). The top 10 genera for the highest performing treatment group are indicated in the legend for each dataset.

To statistically compare overall diversity, we tested for differences between treatment groups using Simpson’s and Shannon’s indices from all three studies (Supplemental Table 3). Among the four treatment group comparisons, none were significantly different for either Simpson’s nor Shannon’s indices (P > 0.05).

### Relative abundance - hypothesis-driven approach

Prior results from the three datasets identified 16 bacterial genera with significant differences in relative abundance in the endurance microbiome (Supplemental Table 2). However, many of the 16 target genera were not found in all four datasets using our bioinformatics pipeline. For example, *Ezakiella* was not identified in any of the datasets (no reads mapped; Supplemental Figures 1-4), although it was confirmed to be present in the BLAST database used to classify sequences in the Geneious workflow. *Ruminiclostridium* and *Actinobacillus* were missing from the Chongqing half-marathon and cyclist datasets. *Methanobrevibacter* and *Pseudobutyrivibrio* were not found in the cyclist dataset and *Akkermansia* was not detected in the Boston marathon dataset.

Among the ten target genera detected in all three datasets, none exhibited significant differential abundance in all four treatment group comparisons (Table 2). Only two genera had significantly different relative abundances in more than one of the treatment group comparisons (*Romboutsia* in the half-marathoners and cyclists, and *Veillonella* in the half-marathoner and marathoner control vs. “athletes after” comparison) (Figure 2). The following sections report results for each dataset individually.

**Figure 2.**
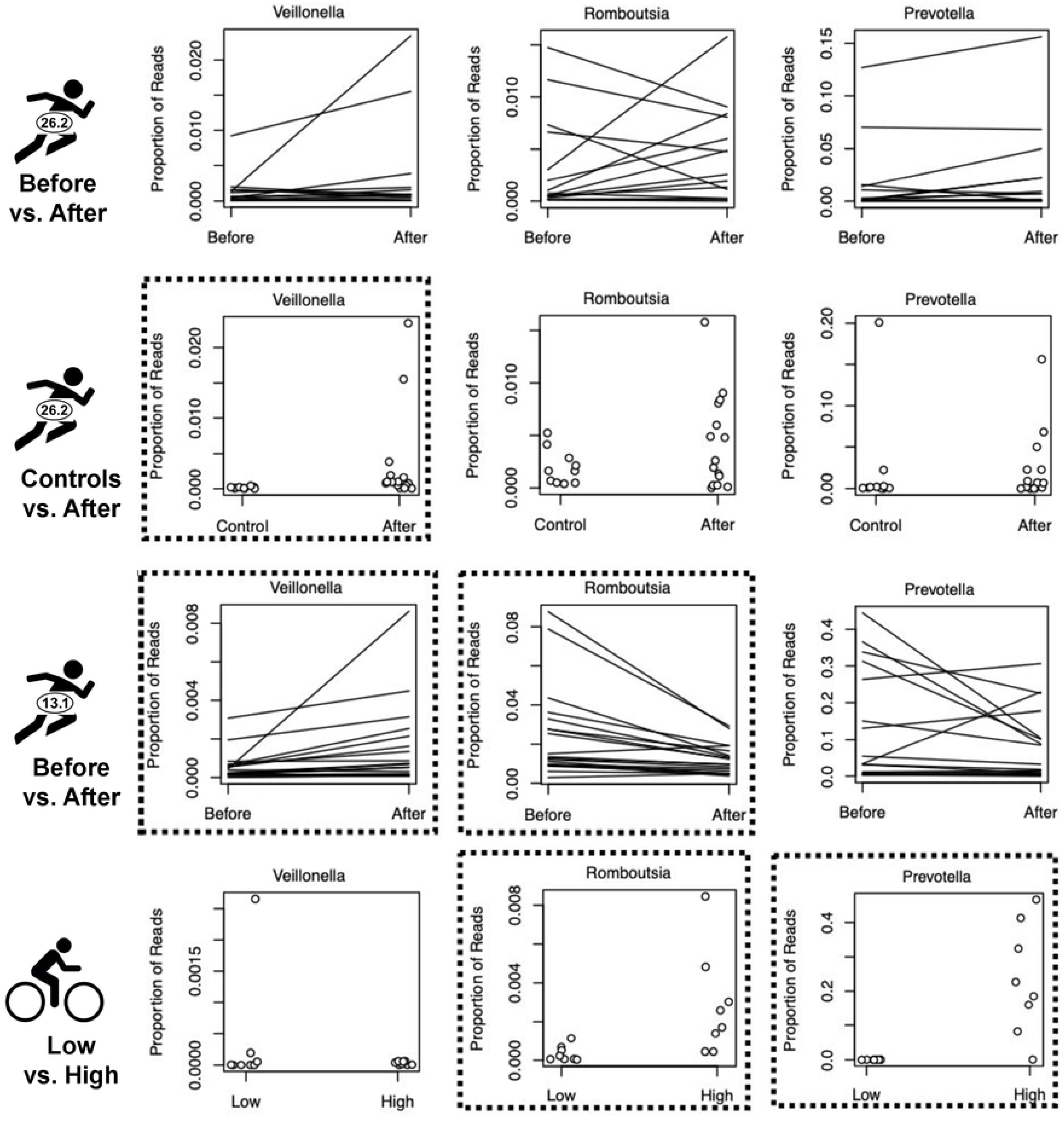
Relative abundance comparisons. Relative abundance for three selected bacterial genera comparing Boston Marathoners before vs. after (A), sedentary controls vs. athletes after (B), Chongqing Half-Marathoners before vs. after (C), and cyclists in the low volume vs. high volume training groups (D). Lines connect paired samples of the same individual (before vs. after), whereas points compare samples taken from different individuals (controls vs. athletes or low vs. high training groups). Significant differences between treatment groups are indicated with black boxes.

**Table 2.**
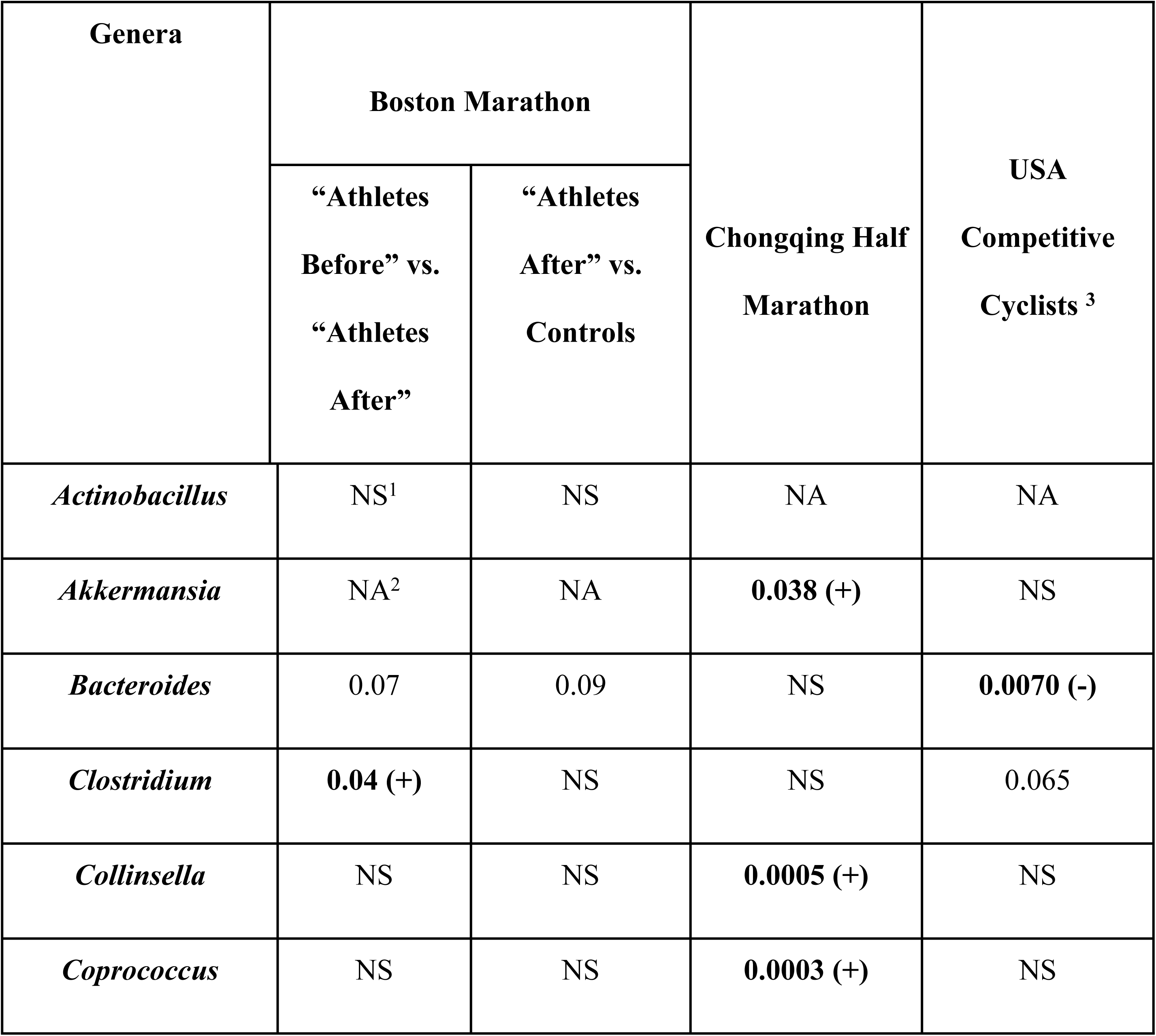
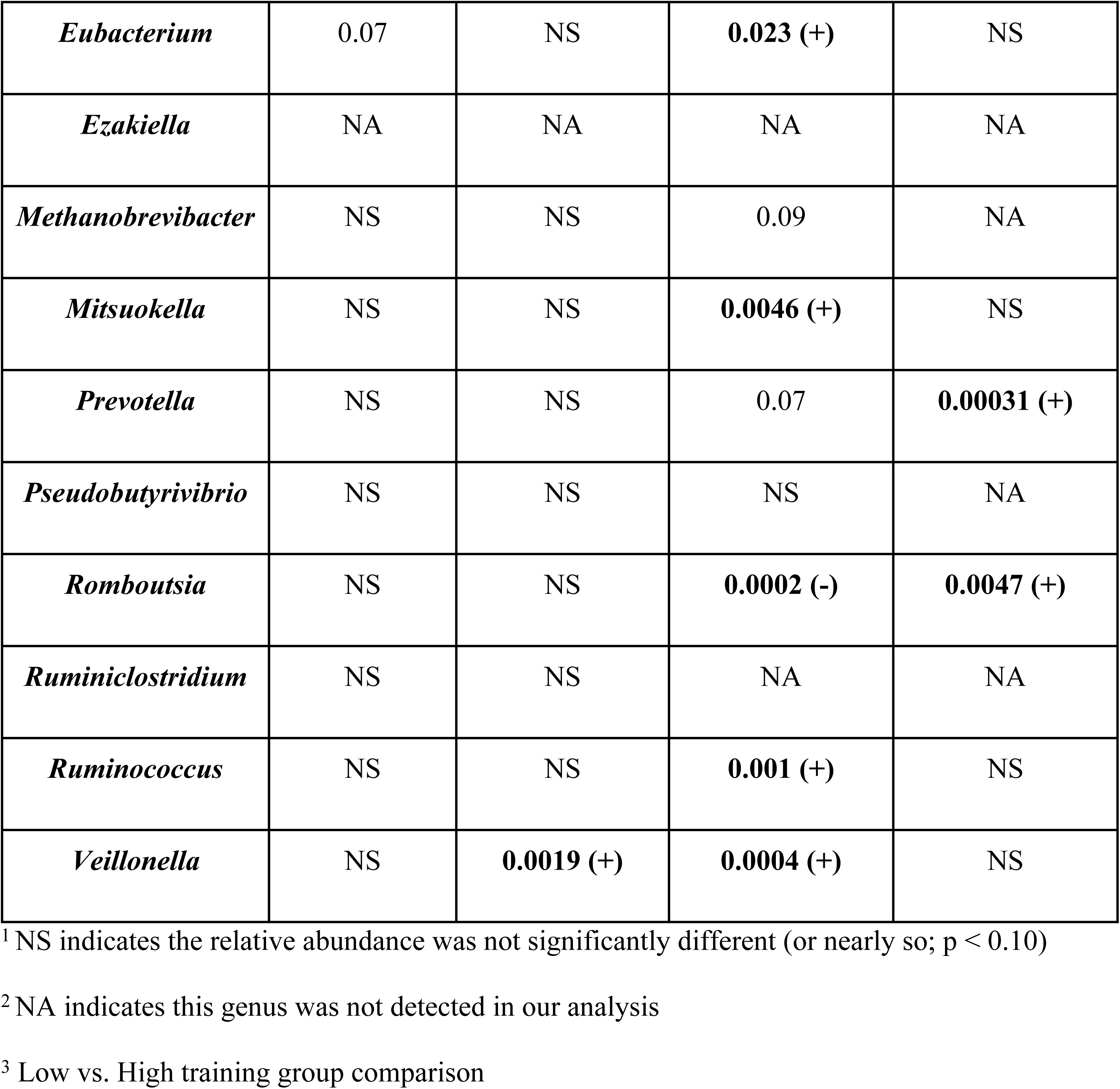
Relative abundance statistical results. P-values from Wilcoxon rank sum tests for the relative abundance comparisons for the 16 target genera among the four treatment group comparisons (Significant results in bold; p < 0.05).

In the Boston marathon dataset, there were two treatment group comparisons and each had one target genus out of the fourteen target genera detected that exhibited significant differences in relative abundance (Figure 2B, Table 2, Supplemental Figures 1 and 2). In comparing “athletes after” to sedentary controls, the average proportion of *Veillonella* in “athletes after” is 22.5-fold larger than in controls (Figure 2A and B, Table 2, Supplemental Table 4, Wilcoxon Test, W = 129, p = 0.002). When comparing “athletes before” vs. “athletes after”, the average proportion of *Clostridium* is 3.6-fold larger in “athletes after” than in "athletes before" (Supplemental Figure 1, Table 2, Supplemental Table 4, Wilcoxon Test, V = 24, p = 0.041).

In the Chongqing half-marathon dataset, eight of the thirteen target genera detected showed significantly different abundances when comparing before and after the half-marathon (Figure 2C, Table 2, Supplemental Figure 3). The four most significant results stand out from the rest (Supplemental Table 2). Three of these four genera showed increased abundance after the event (*Coprococcus, Veillonella,* and *Collinsella*) ranging from 1.97 to 2.67-fold higher (Figure 2C, Supplemental Table 4, Supplemental Figure 3). In particular, *Veillonella* had 2.67-fold higher abundance after the event compared to before the event (Table 2 and Supplemental Table 4; Paired Wilcoxon Test, V = 14, p = 0.0004). Alternatively, *Romboutsia* decreased approximately two-fold after the event (Figure 2, Table 2, Supplemental Table 4; Paired Wilcoxon Test, V = 179, p = 0.0002).

Comparing athletes between the low (n = 8) vs. high (n = 8) training groups from the cyclist dataset, we found three target genera with significantly different abundances (Figure 2D, Table 2, Supplemental Figure 4). *Romboutsia* had approximately 8-fold higher abundance in the high training group compared to the low training group (Figure 2D, Table 2, Supplemental Table 4; Wilcoxon test, W = 6, P = 0.005). *Prevotella* had 700-fold higher abundance in the high training group compared to the low training group (Figure 2D, Table 2, Supplemental Table 4; Wilcoxon test, W = 1, P = 0.0003). *Bacteroides* had 39% lower abundance in the high training group compared to the low training group (Table 2, Supplemental Table 4, Supplemental Figure 4; Wilcoxon test, W = 57, P = 0.007).

### Relative abundance - data exploration (all pairwise comparisons)

After expanding our analyses to compare all genera detected in all four treatment group comparisons, only one genus had significantly different relative abundance after Bonferroni correction (Supplemental Table 4). *Romboutsia* (one of our 16 target genera) from the Chongqing half marathon dataset (see Hypothesis Driven Approach section above), was ∼2x lower after the event compared to before the event (Table 2).

From the Boston marathon dataset, 17 additional genera initially had significantly different abundances (p < 0.05) when comparing sedentary controls vs. “athletes after”, however none of them were significant after the Bonferroni correction (Supplemental Table 4). In comparing Scheiman et al.’s "athletes before" vs. “athletes after”, three additional genera exhibited initially significantly different abundances (*Enterocloster, Fournierella,* and *Marvinbryantia*), however once again, none of them were significant after the Bonferroni correction (Supplemental Table 4). Among the half marathoners, there were 28 additional genera with p < 0.05, but none outside of *Romboutsia* from our 16 target genera were significant after Bonferroni correction. In comparing low vs. high cyclist training groups, we found two additional genera among the non-target genera with initially significant differences (*Turicibacter* and *Bacteroides*, P < 0.01), but neither were significant after Bonferroni correction. *Prevotella* also had P < 0.01, but was one of the target genera, so no Bonferroni correction was necessary (described above).

### Changes in bacterial associations

To determine if the endurance microbiome has distinct associations among bacteria, we first tested for all pairwise correlations in relative abundances between bacterial genera detected across datasets. Second, we used hierarchical clustering to identify networks of bacterial associations to determine whether there are any significant differences among treatment groups.

*Correlated Abundances* **-** First, we tested for significant correlations in all pairwise comparisons of bacterial abundances and then we looked for changes in these associations between treatment groups. Neither approach detected any consistent changes in bacterial associations in endurance microbiomes among all the datasets (Figure 3).

**Figure 3.**
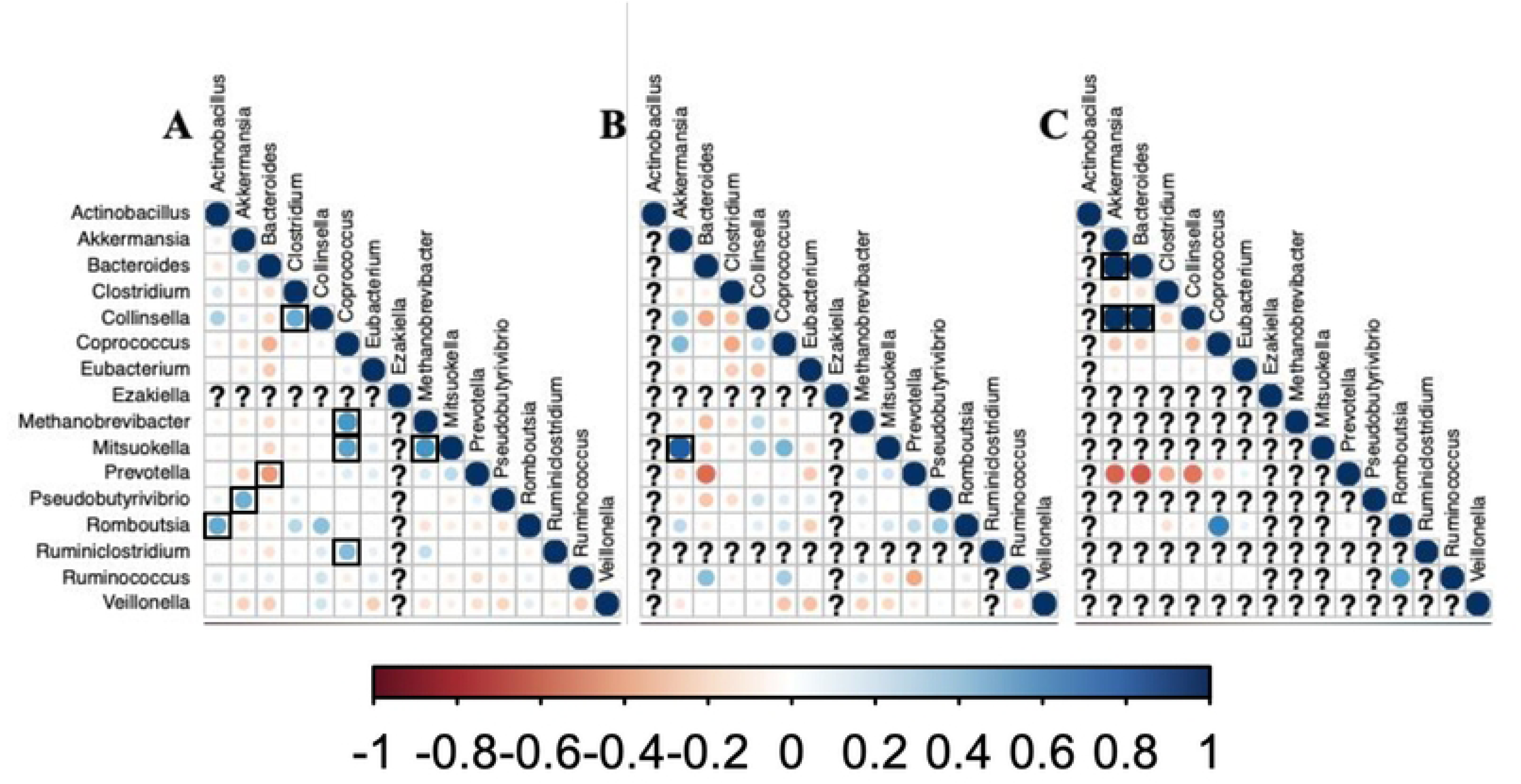
Pairwise correlations between bacterial genera. Pairwise Pearson’s correlations of bacterial genera after the Boston Marathon (A), after the Chongqing Half Marathon (B), and among USA cyclists in the high training group (C). The direction of the correlation (r) is indicated with the color of the circle (red = negative correlation; blue = positive correlation).

Stronger correlations are indicated with greater color intensity and larger diameter of the circle. Significant correlations are boxed in black (p < 0.05 after Bonferroni correction). Question marks represent taxa that were not detected in each dataset.

No correlations in the top 10 most significant correlations after vs. before the endurance event or in high vs low training groups are the same in all four datasets (Table 3, Figure 3). *Blautia* & *Dorea*, *Anaerobutyricum* & *Dorea*, *Anaerobutyricum* & *Streptococcus, and Dorea* & *Streptococcus* are in the top 10 significant correlations in “athletes after” but not in “athletes before” and in “athletes after” but not in controls from the Boston Marathon samples. No other correlations are the same in any two datasets (or more) (Supplemental Table 5). Detailed descriptions of changes in pairwise bacterial associations for each of the individual studies can be found associated with Supplemental Table 5.

**Table 3.** Changes in bacterial correlations. Correlations between bacterial genera that changed from not correlated to strongly correlated (in the top ten most significant correlations) in endurance microbiome comparisons. Target genera are in bold. R-value and P-value for significant correlations are provided.

| Dataset | Taxon 1 | Taxon 2 | R-value | P-value |
| --- | --- | --- | --- | --- |
| **Scheiman et al.**<br>**Athletes After vs**<br>**After Before<sup>1</sup>** | *Blautia* | *Streptococcus* | 0.865 | 0 |
|  | *Blautia* | *Dorea* | 0.859 | 0 |
|  | *Anaerobutyricum* | *Dorea* | 0.853 | 0 |
|  | *Flavonifractor* | *Holdemania* | 0.833 | 0 |
|  | *Anaerobutyricum* | *Streptococcus* | 0.817 | 4.44E-16 |
|  | *Dorea* | *Streptococcus* | 0.792 | 1.07E-14 |
| **Scheiman et al.**<br>**Athletes After vs**<br>**Controls** | *Anaerobutyricum* | *Blautia* | 0.895 | 0 |
|  | *Blautia* | *Dorea* | 0.859 | 0 |
|  | *Anaerobutyricum* | *Dorea* | 0.853 | 0 |
|  | *Butyricicoccus* | *Mediterraneibacter* | 0.840 | 0 |
|  | *Anaerobutyricum* | *Streptococcus* | 0.817 | 4.44E-16 |
|  | *Barnesiella* | *Phascolarctobacterium.* | 0.813 | 4.44E-16 |
|  | *Dorea* | *Streptococcus* | 0.792 | 1.07E-14 |
|  | *Anaerostipes* | *Butyricicoccus* | 0.787 | 2.13E-14 |
| **Zhao et al.<sup>2</sup>** | *Eggerthella* | *Flavonifractor* | 0.969 | 1.01E-11 |
|  | *Eggerthella* | *Hungatella* | 0.959 | 8.91E-11 |
|  | *Desulfovibrio* | *Enterococcus* | 0.949 | 5.85E-10 |
|  | *Eggerthella* | *Erysipelatoclostridium* | 0.943 | 1.51E-09 |
|  | *Intestinibacter* | *Lacrimispora* | 0.942 | 1.66E-09 |
|  | *Faecalimonas* | *Lacrimispora* | 0.941 | 2.14E-09 |
|  | *Klebsiella* | *Streptococcus* | 0.941 | 2.14E-09 |
| **Petersen et al.<sup>3</sup>** | ***Collinsella*** | *Flavonifractor* | 0.998 | 1.55E-08 |
|  | *Flavonifractor* | *Lacrimispora* | 0.998 | 3.04E-08 |
|  | ***Akkermansia*** | ***Bacteroides*** | 0.997 | 4.43E-08 |
|  | *Lacrimispora* | *Pseudoflavonifractor* | 0.997 | 4.77E-08 |
|  | *Flavonifractor* | *Pseudoflavonifractor* | 0.996 | 1.42E-07 |
|  | ***Akkermansia*** | *Flavonifractor* | 0.995 | 2.47E-07 |
|  | ***Collinsella*** | *Lacrimispora* | 0.995 | 3.89E-07 |
|  | ***Collinsella*** | *Ruthenibacterium* | 0.994 | 5.25E-07 |
|  | ***Akkermansia*** | ***Collinsella*** | 0.994 | 5.44E-07 |
|  | *Flavonifractor* | *Ruthenibacterium* | 0.994 | 5.51E-07 |

<sup>1</sup> P < 3.13E-5 is the cutoff for significance after the Bonferroni correction for Scheiman et al.
<sup>2</sup> P < 1.62E-5 is the cutoff for significance after the Bonferroni correction for Zhao et al.
<sup>3</sup> P < 5.807E-05 is the cutoff for significance after the Bonferroni correction for Petersen et al.

*Networks of Bacterial Associations (NetCoMi) -* The bacterial association networks are statistically similar between treatment groups within a dataset and very different across datasets (Supplemental Figures 5-8). First, comparing treatment groups within each dataset, the Rand index (which measures the similarity of two networks by randomly permuting the labels) was always positive and ranged from 0.121 to 0.507. In all cases, Rand was significantly different from zero indicating that the networks within a dataset were more similar to each other than randomly shuffled networks [42]. Additionally, the number of hub taxa was significantly different between treatment groups.

However, the variation across datasets made the networks largely incomparable and no consistent patterns emerged in standard network metrics when comparing across networks (Supplemental Table 6) nor upon visual inspection of target genera. Detailed descriptions of the network results for each of the individual studies can be found associated with Supplemental Table 6.

## Discussion

We reanalyzed the raw data from three gut microbiome datasets of endurance athletes in search of a universal endurance microbiome. By applying the same bioinformatics workflow and downstream statistical analyses across these diverse datasets, we controlled for one source of variation. Overall, we did not detect any hallmarks of a universal endurance microbiome among these datasets. Similar conclusions have recently been reached by Sato and Suzuki [43] who summarized their literature review by writing, “Collectively, our findings suggest that intestinal microbiota diversity is more likely to vary among individuals than to be affected by ultramarathon.”

### Diversity

Alpha diversity can be measured several ways (richness, Simpson’s index, Shannon’s index, etc.). However, in our reanalyses, there were no consistent differences in alpha diversity among the four treatment group comparisons. In direct comparison to the results previously reported from these datasets, Scheiman et al. did not report any diversity statistics [24], Zhao et al. [23] reported no significant differences in alpha diversity after finishing the half-marathon, but did detect a change in diversity using taxonomic profiling [23], and Petersen et al. [18] detected higher diversity in a subset of their data (“Cluster Three”), but not a universal change in bacterial diversity associated with increased training duration [18]. More generally, several other studies have reported increased microbial diversity after an endurance event [6, 16, 19, 20, 44–46] (only when using Simpson’s index for cross country skiers, though). However, there are also numerous investigations that found no change in microbial diversity in endurance athletes [10, 20, 22, 43, 47] (only when using Simpson’s index for marathon runners). Our results and the mixed results from the literature cited above clearly indicate that endurance training does not universally (nor even consistently) increase measures of bacterial diversity in the gut microbiome of endurance athletes.

### Differences in relative abundance

In our reanalyses, none of the 16 target genera exhibited consistently significant differential abundance between treatment groups and across all four studies. The most encouraging results were from two genera that were significantly different in two of the four treatment group comparisons (*Romboutsia* and *Veillonella*). Nine target genera were significant in only one of the comparisons and seven target genera had no reads detected from one or more of the datasets. In fact, one target genus (*Ezakiella*) was not detected in any of the three datasets. (These don’t add to 16 since some genera were missing in one dataset and significant in another dataset, such as *Akkermansia.*)

The two differentially abundant genera found in two studies (*Veillonella* and *Romboutsia*) were from Boston marathoners Athletes After vs. Control and Chongqing half marathoners (*Veillonella*) and Chongqing half marathoners and cyclists (*Romboutsia*), but both findings were curious. First, although we detected significantly higher abundance of *Veillonella* in “athletes after” the Boston Marathon vs. sedentary controls, Scheiman et al. reported *Veillonella* as the sole bacterial taxon differentiating “athletes before” vs. “athletes after” (it was not significant in their “athletes after” vs. controls, but they did note that it was “more prevalent among runners than non-runners”). *Veillonella* abundances exhibit very high levels of variation in “athletes after”, however our non-parametric analysis (Wilcoxon Rank Sum Test) should have accommodated these outliers (as was implemented in [24]). Unlike Scheiman et al., we averaged all samples from “athletes before” and “athletes after”. In contrast Scheiman et al. performed regression against time before and after the marathon - suggesting the bacteria can anticipate the upcoming endurance event, which does not seem realistic (see Fig 1c in [24]).

For the second genus detected in two datasets, Zhao et al. [23] discuss *Romboutsia* in their text, but in their main figure, *Romboutsia* appears to have been replaced with “*Phaseolus sativus*”, a flowering plant in the pea family. Assuming this legume is really *Romboutsia* in disguise, both our analysis and that of Zhao et al. show it decreases in relative abundance after the half-marathon. However, *Romboutsia* abundance increases in the high training group of Petersen et al. (2017) and in our reanalysis thereof. Unless the effects of *Romboutsia* are opposing in cyclists vs. half- marathoners, it appears that even in the two most hopeful hallmarks of a universal endurance microbiome, we find the results are inconsistent, at best.

Outside of these four treatment group comparisons, several studies have compared relative abundance of bacterial taxa in endurance microbiomes (recently reviewed in [48] and [49]). Each study reports a different number and identity of the bacterial genera with differential abundance [10, 16, 19, 20, 22, 43, 46, 47, 50–55]; (also see Supplemental Table 1). Miranda-Comas’ review of the gut microbiome in sports reports, “Although there is great variation in studies, *Faecalibacterium prausnitzii, Roseburia hominis, Akkermansia muciniphila,* and *Prevotella* species are some of the most commonly referenced as healthy or health- promoting gut species” [56]. The inconsistencies in the number and identity of the bacterial genera unique to the endurance microbiome from a wealth of independent studies question its existence.

### Changes in associations

If there were some universality to the endurance microbiome, it may exhibit consistent associations among bacteria due to the unique gut environment of endurance athletes. Numerous studies have found a negative correlation between *Prevotella* and *Bacteroides* [18, 10, 20, 16, 26, 57, 58]. We used this established relationship as a positive control to confirm our analyses were accurate. For the Boston Marathon dataset, over all three treatment groups, *Prevotella* and *Bacteroides* were negatively correlated (r = -0.38; p = 5.6E-8) and ranked 86 out of 152 significant correlations overall. In the Chongqing half marathon dataset, *Prevotella* and *Bacteroides* have the strongest negative correlation among the target genera, yet this relationship was not significant after Bonferroni correction (r = -0.48; p = 0.002). In the cyclists dataset, the strongest negative correlation was between *Bacteroides* and *Prevotella* (r = -0.59, p = 0.0003). The negative correlation between these two cosmopolitan genera provide an internal control on our method, but are unlikely related to the endurance microbiome as this relationship is reported in a range of gut microbiome studies.

After establishing the negative relationship between *Prevotella* and *Bacteroides*, we investigated all pairwise correlations among gut bacteria. Although each of the four treatment group comparisons identified several pairs of genera, each dataset had different genera emerging with significant associations. We found three treatment group comparisons with none of the 16 target genera involved in the top emerging correlations (both marathon treatment group comparisons and the half marathon dataset). One dataset (cyclists Low vs. High training groups) had three of the 16 target genera. Furthermore, only one genus, *Streptococcus* (correlated with *Klebsiella*), appears in the top 10 changing correlations of more than one dataset (marathoners and half marathoners). Clearly associations between these target genera (and the rest of the microbiome for that matter) are not changing in a predictable, consistent manner in endurance athletes.

Matchado et al. warn that “…it is challenging to differentiate between direct and indirect associations, in particular if these are related to environmental factors” and “microbial association networks are typically undirected and not all interactions represent true ecological relationships” [35]. However, in microbial networks, “hubs help pinpoint the most important taxa of a given environment, thereby providing essential clues about how specific taxa or gene products may contribute to ecosystem functioning” [41]. Similar criticisms about the meaning of correlated abundances and networks can be found in the ecological literature [59]. However, detecting correlations and identifying associations can be helpful inroads into the dynamics of the gut microbial community, which can become hypotheses for more manipulative tests under controlled and experimental conditions to determine the underlying causes of the correlations (i.e. facilitation, competition, use of a shared resources, etc.). Unfortunately, we found only vast differences among datasets when examining networks of bacterial genera in endurance athletes’ gut microbiomes.

Are all microbiome bacteria in everyone, just needing the right environment to thrive, or is the microbiome constrained by the available bacteria? Restated, “is the endurance microbiome growth-limited or colonization-limited”? Truly answering this requires better understanding the sensitivity and limits of detection of current approaches to microbiome analysis. Scheiman et al. [24] blur the boundaries between the two hypotheses in writing that “An important question is how this performance-facilitating organism first came to be more prevalent among athletes. We propose that the high-lactate environment of the athlete provides a selective advantage for colonization by lactate-metabolizing organisms such as *Veillonella*.” Some degree of colonization limitation could logically encourage the use of probiotics (e.g., “Nella” Probiotics). Our lack of consistent correlations among datasets using Pearson’s pairwise correlations and the network approaches, and repeated references to the large inter-individual variation in gut microbiomes in general, also suggests that gut microbiomes of endurance athletes are colonization-limited, and that most bacteria are not omnipresent in the gut awaiting the right environmental cues. Whether introducing target taxa like *Veillonella* or *Romboutsia* into the gut microbiomes of endurance athletes with conducive environments for colonization has a consistent response will require much broader sampling (but see Scheiman et al. 2019 extension to germ-free mice).

The excess of positive correlations compared to negative correlations detected in all datasets is likely a technical limitation [60, 61]. Although one might be tempted to infer a higher frequency of commensal relationships compared to competitive interactions, Badri et al. explains that, “the positive skewness may also be due to technical limitations in the data generation process and shortcomings in current statistical estimation. For instance, truncation to zero effects for low sequencing read counts likely obstructs unbiased estimation of negative correlations” [36].

### Limitations

Early gut microbiome studies classified most samples into one of a few narrow enterotypes, especially with regard to the frequency of *Prevotella* and/or *Bacteroides.* However, thorough meta-analysis clearly showed that these enterotypes were not reflective of the data [26, 62]. Similar to the passing of the enterotypes concept, our reanalysis of these datasets clearly indicate that there is no universal gut microbial community shared by endurance athletes.

Our results have several limitations, some of which are technical, and others biological. One disconcerting practical limitation was that many of the published studies we sought to follow up and reanalyze did not have publicly available 16S rRNA sequence data for reanalysis via the pipeline developed for this study (Supplemental Table 1).

Another technical limitation surrounds the use of putatively “universal” 16S primers. Several authors have pointed out that, although the use of 16S metagenomics for determining microbial composition of the gut is powerful [63], the standard 16S primers may not amplify all bacteria in equal proportions to their starting population sizes [64]. However, the three studies and datasets that we reanalyzed used the same primers, so any errors would have affected all datasets equally. This issue is therefore unlikely to explain our failure to detect a universal endurance microbiome. However, because we narrowed our focus to genera for which more than 75% of samples had reads mapped, it is possible that microbes whose rRNA sequences are poorly amplified by these universal primers would have been missed.

A more biological limitation of our study is the limited taxonomic granularity we were able to apply. We narrowed our master taxa list to include only taxa with genus-level taxa identification, but it is well known that different microbial species within a genus may have distinct impacts on human metabolism and physiology (e.g. *Clostridium*). We may therefore have missed species-level or even within species strain-level changes or interactions relevant to endurance athletes. More broadly, a 16S profiling approach will be largely blind to changes in microbial populations related to mobile genetic elements such as plasmids, which can significantly impact microbial physiology [65].

Generally, because there are so many factors that can affect the microbiome, any comparison of microbiomes must be sufficiently controlled and have adequate sample size to account for these factors. Diet, which can be highly variable and is often not well controlled in research studies with humans, is undoubtedly one of the largest factors affecting the gut microbiome. Challenges related to diet were clearly evident in our analysis. The study of Chinese half-marathoners examined here showed a microbiome response in just one day to the chemistry of the dietary intake. Clarke et al. found that athletes had higher protein intake, which may have affected microbial diversity in athletes’ gut microbiomes, regardless of their endurance-induced physiological differences [16].

Indeed, life-long effects of early diet on the gut microbiome have been repeatedly reported [26, 66], that could constrain future exercise-induced changes. In addition to diet there are dozens of other environmental factors that strongly impact the composition of the gut microbiome. A rule of thumb for a logistic regression is 10-20 samples per parameter being estimated [67] which suggests on the order of 100-200 samples to account for the numerous covariates affecting the gut microbiome. The number of covariates can be reduced (but not eliminated) by using a paired sampling scheme (i.e. the same athlete before vs. after), however, this approach fails to detect long-term changes in gut microbiomes that differentiate endurance athletes from sedentary controls.

Finally, we conducted a power analysis to estimate the sample size necessary to detect significantly different abundances between treatment groups. We used the empirically determined d based on the ability to detect a 10% difference in the average abundance per genus and calculated the pooled standard deviation among all samples per genus. We applied a power of 0.8 (an 80% chance of concluding there’s a real effect) and a Bonferroni corrected alpha based on the number of genera detected per dataset assuming that many tests will be conducted. Our power analysis suggests that all four treatment group comparisons require substantially larger sample sizes than those in the published studies, ranging from ∼150-fold more samples (Chongqing half marathon) to ∼800-fold more samples being necessary (Boston marathon Athletes After vs Controls) (Supplemental Table 7). Even using the largest d value among all genera detected per study, the recommended sample sizes are still 10-fold to 150-fold smaller than necessary. Ours and other meta-analyses have combined studies of different sports (e.g. O’Donovan et al. 2020 [46] examined athletes across 16 sports), but in the end we believe that this approach cannot substitute for what is really needed to understand the endurance microbiome (if it exists): well-controlled studies that generate orders of magnitude more data from large sample sizes. Only then will investigators be able to reliably detect significant effects of intensive exercise on the gut microbiome against a background of confounding covariates.

### Future research

Rather than a “universal” endurance microbiome, it must be considered that different types of athletic activity may place different demands on the anatomy and physiology of participants, that then differentially influence the gut microbiome. Tabone et al. [22] write that “Exercise frequency, intensity, performing time, type of exercise, exercise volume and progression are all factors that influence physiological responses and exercise adaptations, and will need to be considered in future studies investigating the beneficial effect of exercise on the gut microbiota” [22]. Due to inter-individual variation, more ideally controlled studies could compare the same individual sampled repeatedly before and after endurance events. Sedentary control groups could be thoroughly sampled, under diet controlled conditions, then turned into an endurance group and sampled during the process. Pugh et al. [14] recently reviewed the differences between the gastrointestinal health of female endurance athletes remarking, “The links between female microbiome, estrogen, and systemic physiological and biological processes are yet to be fully elucidated” and that “Many of the male-female differences seen (e.g. in immune function) may be, at least in part, influenced by such GI related differences” [68]. Given these complexities, our search for the existence of a universal endurance microbiome may be destined to mirror the mixed results of probiotic use in sports [13] - elusive at best, and non-existent at worst.

## Data availability statement

All read counts and proportions from our Geneious workflow are available from Figshare (DOI: 10.6084/m9.figshare.c.6036347). We have also included all the R scripts used for statistical analyses and our Geneious workflow that can be downloaded and run in Geneious Prime 2021.2.2.

## Acknowledgments

We thank the SCU Department of Biology staff for computational support and SCU’s undergraduate research initiatives, specifically the DeNardo Scholar program (KT) and REAL program (HO) for financial support for undergraduate researchers. The Santa Cruz Track Club provided an opportunity to present our initial findings to their membership in a virtual seminar. Dr. Peter Turnbaugh (UCSF) and Dr. Tracy Ruscetti (SCU) shared critical advice at an early (pre-pandemic) stage in this study.

## Supporting information

**Supplemental Table 1**. Dataset characteristics for studies that were considered for reanalysis. The first three criteria had to be satisfied as well as one or both of the last two criteria. Datasets chosen for reanalysis are indicated in bold.

**Supplemental Table 2**. Source of 16 target genera and statistical results from previous studies examined in this work.

**Supplemental Table 3**. Simpson and Shannon diversity indices among studies and treatment groups therein. None of the comparisons between treatment groups were significantly different at alpha = 0.05.

**Supplemental Table 4**. Significant differences in relative abundance based on Wilcoxon tests for all genera (data exploration) from Scheiman’s “Athletes Before” vs. “Athletes After”, Scheiman’s Controls vs. “Athletes After”, Zhao’s “Athletes Before” vs. “Athletes After”, and Petersen’s low vs. high training groups. Genera are separated by dataset, then sorted by increasing p-value.

**Supplemental Table 5**. Significant correlations per treatment group between all pairwise bacterial genera combinations sorted by decreasing r-value. (discussion of these individual results follows)

**Supplemental Table 6**. Network descriptors for bacterial community associations among the top 100 bacterial genera in each treatment group for the three datasets. LCC = largest connected component; Dissimilarity = 1 - edge weight

**Supplemental Table 7**. Power analysis of four treatment group comparisons reanalyzed herein to determine the recommended sample size.

Supplemental Figure 1. Relative abundance comparisons for Scheiman et al. “athletes before” vs. “athletes after” the Boston Marathon for the sixteen target genera previously reported to be part of the endurance microbiome. Significant differences between treatment groups are indicated with black boxes. NA indicates the genus was not detected in the microbiome. Lines connect means of multiple samples per individual.

Supplemental Figure 2. Relative abundance comparisons for Scheiman et al. controls vs. “athletes after” the Boston Marathon for the sixteen target genera previously reported to be part of the endurance microbiome. Significant differences between treatment groups are indicated with black boxes. NA indicates the genus was not detected in the microbiome. Points on the x-axis were jittered to increase visibility. Points represent means of multiple samples per individual.

Supplemental Figure 3. Relative abundance comparisons for Zhao et al. “athletes before” vs. “athletes after” the Chongqing International Half-Marathon for the sixteen target genera previously reported to be part of the endurance microbiome. Significant differences between treatment groups are indicated with black boxes. NA indicates the genus was not detected in the microbiome. Lines connect the same individual sampled at two timepoints.

Supplemental Figure 4. Relative abundance comparisons for Petersen et al. USA cyclists high vs. low training groups for the sixteen target genera previously reported to be part of the endurance microbiome. Significant differences between treatment groups are indicated with black boxes. NA indicates the genus was not detected in the microbiome. Points on the x-axis were jittered to increase visibility.

Supplemental Figure 5. Network comparison for Scheiman et al. comparing “athletes before” and “athletes after” the Boston Marathon. Nodes are bacterial genera. Node colors indicate clusters, line colors indicate positive associations (green) and negative associations (red), line weights reflect eigenvalues (connectedness). Line lengths are arbitrary. Hubs (bold font) are nodes with an eigenvector centrality above the empirical 95% quantile of all eigenvector centrality values.

Supplemental Figure 6. Network comparison for Scheiman et al. comparing sedentary controls to “athletes after” the Boston Marathon. Nodes are bacterial genera. Node colors indicate clusters, line colors indicate positive associations (green) and negative associations (red), line weights reflect eigenvalues (connectedness). Line lengths are arbitrary. Hubs (bold font) are nodes with an eigenvector centrality above the empirical 95% quantile of all eigenvector centrality values.

Supplemental Figure 7. Network comparison for Zhao et al. comparing athletes before to athletes after the Chongqing International half-marathon. Nodes are bacterial genera. Node colors indicate clusters, line colors indicate positive associations (green) and negative associations (red), line weights reflect eigenvalues (connectedness). Line lengths are arbitrary. Hubs (bold font) are nodes with an eigenvector centrality above the empirical 95% quantile of all eigenvector centrality values.

Supplemental Figure 8. Network comparison for Petersen et al. comparing the low training group of professional cyclists to the high training group of professional cyclists. Nodes are bacterial genera. Node colors indicate clusters, line colors indicate positive associations (green) and negative associations (red), line weights reflect eigenvalues (connectedness). Line lengths are arbitrary. Hubs (bold font) are nodes with an eigenvector centrality above the empirical 95% quantile of all eigenvector centrality values.

Supplemental Files (available in FigShare, DOI: 10.6084/m9.figshare.19799203)

**Supplemental Fi1e 1**. R script and input data to generate diversity index values and compare statistically.

**Supplemental Fi1e 2**. Bioinformatics pipeline (Geneious workflow)

